# Elevated fear responses to threatening cues in rats with early life stress is associated with greater excitability and loss of gamma oscillations in ventral-medial prefrontal cortex

**DOI:** 10.1101/2021.07.28.454150

**Authors:** Florencia M Bercum, Maria J Navarro Gomez, Michael P Saddoris

## Abstract

Stress experienced early in development can have profound influences on developmental trajectories and ultimately behaviors in adulthood. Potent stressors during brain maturation can profoundly disrupt prefrontal cortical areas in particular, which can set the stage for prefrontal-dependent alterations in fear regulation and risk of drug abuse in adulthood. Despite these observations, few studies have investigated *in vivo* signaling in prefrontal signals in animals with a history of early life stress (ELS). Here, rats with ELS experienced during the first post-natal week were then tested on a conditioned suppression paradigm during adulthood. During conditioned suppression, electrophysiological recordings were made in the ventral medial prefrontal cortex (vmPFC) during presentations of a fear-associated cues that resolved both single-unit activity and local field potentials (LFPs). Relative to unstressed controls, ELS-experienced rats showed greater fear-related suppression of lever pressing. During presentations of the fear-associated cue (CS+), neurons in the vmPFC of ELS animals showed a significant increase in the probability of excitatory encoding relative to controls, and excitatory phasic responses in the ELS animals were reliably of higher magnitude than Controls. In contrast, vmPFC neurons in ELS subjects better discriminated between the shock-associated CS+ and the neutral (“safe”) CS-cue than Controls. LFPs recorded in the same locations revealed that high gamma band (65-95 Hz) oscillations were strongly potentiated in Controls during presentation of the fear-associated CS+ cue, but this potentiation was abolished in ELS subjects. Notably, no other LFP spectra differed between ELS and Controls for either the CS+ or CS-. Collectively, these data suggest that ELS experience alters the neurobehavioral functions of PFC in adulthood that are critical for processing fear regulation. As such, these alterations may also provide insight into to increased susceptibility to other PFC-dependent processes such as risk-based choice, motivation, and regulation of drug use and relapse in ELS populations.

## Introduction

Early Life Stress (ELS) has been identified as a particularly potent risk factor due to the insults and stressors occurring during critical developmental time windows in brain maturation. In human populations, ELS is a result of neglect, abuse and/or trauma experienced before the age of 18. Child Protective Services investigated 3.5 million cases of possible child maltreatment in 2018, an 8% increase from 2014 in the United States. However, estimating the prevalence of ELS is difficult as most cases are unreported. Human and animal studies refer to perinatal stress as exposure to acute and/or chronic stressors during prenatal and early postnatal life. While the definition is broad, it aims to highlight a critical period of growth, organogenesis and brain development in which the fetus or child are highly vulnerable to insult. During this period, exposure to stress during early life is associated with higher rates of chronic illness such as diabetes, obesity, cardiovascular, gastrointestinal and respiratory illness as well as autoimmune disorders (McEwen, 2003; Taylor, 2010).

In addition to these somatic responses, ELS can produce profound consequences on the developing central and peripheral nervous system (Lupien et al., 2009; McEwen, 2003). The prevalence of ELS in mental illness is alarmingly high such that 50% to 64% of patients diagnosed with depression, anxiety or substance use disorders report exposure to early life adversity (Dube et al., 2003; Enoch, 2011; Vogt et al., 2016). Clinical studies have also reported a positive correlation between the intensity and duration of ELS and the number of psychopathologies an individual develops, as well as symptom severity (Carr et al., 2013). Indeed, a host of stress and anxiety disorders such as generalized anxiety disorder, panic disorder, post-traumatic stress disorder (PTSD) and obsessive-compulsive disorder have the highest rate of co-occurrence and are comorbid with substance use disorders (SUD) (Regier et al., 1990), all of which have been linked to ELS as a predictor of their development (Brady and Sinha, 2005; Brown and Barlow, 1992; Enoch, 2011; McEwen, 2003; Regier et al., 1990).

The rodent and human brain follow a highly-conserved sequences of structural development (Rice & Barone, 2000). Rodents exposed to ELS have reported morphological, neurochemical and behavioral alterations that parallel some of the findings reported in humans (McEwen, 2003; Weinstock, 2017, 2008). Unlike subcortical structures, in which early cellular processes such as migration, differentiation, synaptogenesis and gliogenesis are well underway, limbic-connected structures such as hippocampus, amygdala and prefrontal cortex (PFC) are in their early stages of development at birth (Herlenius and Lagercrantz, 2004; VanTieghem and Tottenham, 2018). More importantly, the PFC is the last cortical structure to fully develop as it continues these processes into adulthood making the PFC highly susceptible to environmental insult throughout childhood (Kroon et al., 2019; VanTieghem and Tottenham, 2018). During the early stages of development, cortical and subcortical projections reach the PFC to ensure proper communication between the PFC and the rest of the brain, many of which introduce neuromodulatory activity like dopamine into the region (Kalsbeek et al., 1988). These neuromodulators play critical roles in the development of PFC circuits during the first week of postnatal life. Additionally, increased glutamatergic activity during early development dominates cortical pyramidal cell transmission and is critical in synaptogenesis as well. NMDA activity is predominant and aids in fine tuning synapses and in areas of excessive activity it enables apoptosis (Herlenius and Lagercrantz, 2004; Rice and Barone, 2000).

In rodents, the PFC along the medial aspect frontal regions is typically subdivided into two functional divisions, the dorsal medial aspect (*dmPFC*; which includes the prelimbic [PL] cortex) and the ventral medial aspect (*vmPFC*; including the infralimbic [IL] cortex). Though these PFC regions lack direct homology with primate regions (Laubach et al., 2018), evidence exists that functional and developmental overlap between rodent vmPFC and similar human prefrontal regions such as Brodmann’s area 25. For example, this region has been implicated in cognitive and behavioral flexibility deficits in patients that suffer from Anxiety Disorders, ASD and SUD (Greenberg et al., 2013; Jackson et al., 2016; Myers-Schulz and Koenigs, 2012), as well as decreased activity during fear generalization tasks (Greenberg et al., 2013). Similar activity patterns were also detected in individuals suffering from PTSD when presented with trauma associated cues (Shin et al., 2004) and extinction recall (Milad et al., 2009).

In rodents, vmPFC participates in similar functions (Giustino and Maren, 2015). Via connections with limbic-associated targets such as the amygdala complex, the IL is involved in the consolidation, maintenance and expression of extinction learning as well as habitual behaviors (Quirk and Mueller, 2008). Within the domain of fear conditioning specifically, IL activity is both necessary and sufficient to support fear extinction. Stimulation of IL accelerates fear extinction (Adhikari et al., 2015; Bukalo et al., 2021; Milad and Quirk, 2002) and suppresses spontaneous recovery of fear (Kim et al., 2010), while neurons in this area increase activity during the early stages of extinction learning, with cue-elicited phasic activity emerging only after extinction learning has occurred (Sierra-Mercado et al., 2011a). Conversely, pharmacological or optical silencing of this IL pathway in fear extinction learning results in increased freezing behavior and reduced extinction rates (Adhikari et al., 2015; Gutman et al., 2017; Laurent and Westbrook, 2009).

However, ELS appears to alter these normal fear and anxiety-related processes, disrupting conditioned fear as well as decreasing exploration of open arms in an elevated plus maze (Nisar et al., 2019; Oldham Green et al., 2021; Toda et al., 2014). While recent work has provided tremendous *ex vivo* insight into the genetic, epigenetic, and molecular bases for these differences particularly in the PFC (Oldham Green et al., 2021; Torres-Berrío et al., 2019), surprisingly little is known about how mature *in vivo* neurons in ELS-experienced animals encode information about threat and safety during behavior. Using limited experience with wet bedding as variant of a limited bedding model of ELS (Léonhardt et al., 2007; Molet et al., 2014; Walker et al., 2017), we assessed behavioral and *in vivo* neural activity in vmPFC in adulthood of these ELS subjects (along with unstressed Controls) in a conditioned suppression task. Conditioned suppression is a higher-order assay where rats must use the learned value of a threat-associated cue to guide conflicting actions: (1) continue to seek reward despite threats, or (2) engage in defensive behaviors (e.g. freeze immobility) despite the opportunity to earn rewards. Here we report that ELS animals showed more persistent fear-suppressed motivation than Controls, and that during this task, vmPFC neurons displayed greater excitatory responding to fear cues, but decreased high gamma oscillations in the local field potential band.

## Methods

### Subjects

Subjects for this experiment were initially 26 male and female Long-Evans rats (9 male and 17 female) at the start of the experiment. All animals were bred in-house in the CU Boulder vivarium in an AAALAC-approved facility. Subjects were bred against a TH::Cre background (males were TH::Cre-positive Long-Evans [original line sourced from Rat Resource & Research Center (RRRC)] mated with female standard non-transgenic Long-Evans [Envigo]), though we did not use manipulations to selectively target TH-containing cells in this study, so cre status was not assessed in this study in group assignments. TH::cre rats do not differ in behavior from littermate controls, and are routinely used without manipulation as equivalent to cre-negative littermate (Ferland et al., 2019). All rats were bred in-house and maintained on a 12:12 light/dark cycle (lights on 7:00 A.M.–7:00 P.M.) and all testing occurred during the light phase.

The behavioral procedures occurred in two distinct periods. During the first (“early life”) phase of the experiment (PND 0 - PND 21), dams were allowed *ad libitum* access to enriched breeder chow (Teklad) and water in the home cage, while pups were allowed *ad libitum* access to nursing. During the second (“adulthood”) period for the original pups, rats were first tested under ad libitum conditions (PND180), and afterwards (approximately PND 180-PND 320) restricted to 95% of their free feed weight receiving 10-15g of standard laboratory chow (Teklad) provided directly in their home cage after daily test sessions.

During the pre-weaning period (PND 0-PND 21), the dam and pups were housed in a plastic container (48cm (l) X 26cm (w) X 20cm (h)) with approximately 2cm of wood shaving bedding and a laboratory paper twists for enrichment and nesting material. For the remainder of the experiment the home cage consisted of a plastic container (48cm (l) X 26cm (w) X 20cm (h)) with approximately 2cm of bedding. All procedures were performed in accordance with University of Colorado Institutional Animal Care and Use Committee guidelines for the humane use of laboratory rats in biological research.

### Behavior

#### Early life stress (wet bedding and nesting material)

Male and females were paired, and after given time for mating, pregnant females were isolated to gestate without males present. The early life stressor in this study was an incidental water supply malfunction (Hydropac Alternative Watering System), in which one of the two hydropacs on the cage lid ruptured, resulting in exposure to wet and cool bedding (2 cm) and nesting material for the dam and pups. Note that because cages were provided with two hydropacs, the primary stressor here was related to potential thermal loss in the pups and dam, limited *quality* (dry) bedding to build warm “domed” nests, and potential maternal stress related to these conditions. Indeed, prior models of stress experienced by the dam via exposure to wet-bedding for as little as 10h has been shown to alter concurrent maternal behavior for over a week after exposure (Léonhardt et al., 2007). Importantly, in the present study, the wet bedding stressor was not sufficient to cause any loss of individual pups or litters.

Animals were observed on PND1 and PND2 (Thu/Fri) by a lab research staff member, and then ∼48h later (PND4; Sun) when the wet bedding was discovered. At this point, animals were immediately removed from the wet bedding and placed in normal (dry) bedding. During this time in the vivarium, it was our intention to decrease dam stress by limiting human (care staff) interactions in the vivarium in the first few days post-delivery to a minimum. This activity nevertheless included rapid checks that dams had at least one intact hydropac and access to food. As such, dams retained access to *ad libitum* water and food during this wet-bedding period.

We had initially intended the offspring in this study to be used in our lab to establish future breeding lines for our transgenic colony. Because early life stress is known to potentially induce epigenetic changes in individuals (Torres-Berrío et al., 2019), transgenerational transmission of these change could present a confound, and therefore these animals were not considered viable for future breeding. To avoid euthanasia for these subjects, we instead wondered whether this incidental exposure (while unintended) could provide an opportunity to compare how this developmental stressor during the first week postnatal could alter developmental trajectories, particularly in contrast to controls born and reared in otherwise identical conditions. Indeed, unanticipated stressors can provide valuable insights as to how these events affect brain and development (Dallman et al., 1999), and the lead author on this project has an established record of characterizing perinatal models of stress and development in rats (Bercum et al., 2015). Importantly, as this wet bedding exposure was incidental, we immediately discussed with and obtained approval from our veterinarian and IACUC committee before continuing with these studies. With their support and approval, we designated pups who experienced the wet/cold cage the Early Life Stress (ELS) group, while pups reared under identical conditions in the vivarium at the same time but were raised under normal dry bedding and nesting conditions throughout development comprised the Control group. Animals were pair-housed during the post-weaning period into adulthood, and were only separated following intracranial implants just prior to Conditioned Suppression training.

#### Early Life: Juvenile Social Exploration

Social exploration tests were conducted at PND 180 as described previously (Christianson et al., 2011). For this test, each subject was placed in plastic tub cage (48cm X 26cm X 20cm) that contained 2 cm of fresh bedding located in a designated testing room for a 1-hour habituation period. A novel Long-Evans juvenile rat (target) was then introduced into the cage for 3 min. The behavioral response of the subject animals (ELS/Controls) to the target juvenile was recorded with a video camera placed above the cage. The behavior was then assessed by measuring total duration and frequency of social exploration. All videos were scored in a blinded and randomized manner.

#### Adulthood: Conditioned Suppression

As adults, rats were trained in a Conditioned Suppression paradigm. This task consisted of a sequence of three phases: instrumental conditioning [Context A], auditory fear conditioning [Context B], and finally, fear-conditioned suppression test [Context A]. The specific approaches are explained in detail below.

### Instrumental Conditioning

Lever press was established to assess motivation to seek rewards. On the first day of training, rats (n=16) were introduced to a large operant chamber (Context A: 60cm (w) X 56cm (l) X 36cm (h), smooth Plexiglas floors; MED Associates) where they were first magazine trained to obtain food (45mg raspberry-flavored grain pellets, Purina Test Diet) randomly delivered to a centrally-located foodcup on average about every 60s schedule.

On the subsequent days, a lever was extended and rats could press the lever and obtain food. Each session ended when the rat pressed the lever enough times to deliver 25 pellets or 30min, whichever occurred first. If a rat received the 25 pellets within the 30min session limit, on the following day, the subject was promoted to the next reinforcement schedule. Rats were first trained on an FR1 schedule of reinforcement (each press produces a pellet), followed by variable interval schedules of VI-5 (i.e., the first press in each 5-sec bin reinforced), then VI-15, VI-30 and finally VI-60. Following completion of the VI-60 schedule, rats were implanted with bilateral electrophysiological arrays (see below), then allowed at least 7d to recover. Following recovery, rats were retested in the VI-60 schedule to re-establish stable pressing behavior. For rats who failed to perform adequately at the VI-60 schedule, retraining at denser reinforcement schedules until they could once again advance to stable completion of the VI-60 schedule over three consecutive days.

### Fear Conditioning

On the day following completion of the last VI-60 schedule of pressing, rats were trained in a standard tone-shock fear conditioning paradigm consisted of a single 49-min session. In this task, subjects were tested in a novel chamber (Context B: 43cm X 43cm X 53cm, stainless steel grid floor; MED Associates) that was located in a different location in the research facility from the original Instrumental Context A. The first 5 min of the session consisted of a habituation phase where no cues were presented followed by a randomized presentation of a total of 14 trials with a 180s ITI (7 CS+ trials: 30s tone (5000Hz, 80dB) co-terminating with a 0.8mA footshock delivered through the stainless floor grid bars, and 7 CS-trials: 30s tone (3000 Hz, 80dB) presented without any programmed consequences). Average freezing during the cue presentations and inter-trial intervals (ITI) were automatically scored using MED-Associates software.

### Conditioned Suppression

The day after Fear Conditioning training, test subjects were returned to the original Instrumental Context A chamber. As in prior instrumental testing sessions, a lever was presented and presses were reinforced with 45mg pellets on a VI-60 schedule. However in the Conditioned Suppression sessions, after an initial 5min habituation phase where no cues were presented, rats received random presentations of either the previously shock-predictive CS+ cue (30s high tone; 5000 Hz, 80dB; n=7 presentations) or the neutral CS-cue (30s low tone; 3000Hz, 80dB; n=8 presentations). Note that in this Context A, auditory cues were presented without shock under extinction conditions. Cues were presented with an ISI on average of 150s throughout the session. The number of lever presses during baseline period (no cues BL) was compared to pressing during both the CS+ and CS-period to assess the degree of cue-elicited suppression of instrumental activity. Rats received three consecutive Conditioned Suppression sessions to assess the rate of recovery of instrumental pressing with extinction of the fear response.

### Electrophysiological Recording

Single-unit recordings were acquired during Conditioned Suppression test sessions. Using Plexon Omniplex systems (Plexon, Dallas TX) with a sampling rate of 40 kHz, analog voltages recorded at the site of wires relative to a ground wire were amplified with a unity gain head stage. The other connector on the tethered headstage was connected to a 16-channel electrical commutator (Crist Instruments, Hagerstown, MD) to allow free movement in the chamber. Signals from the commutator were then passed to an A/D converter (MiniDigiAmp) where analog voltages were digitized and filtered to capture spikes and local field potentials (LFPs) on dedicated channels for each wire. OmniPlex received synchronized TTL inputs from the MED Associates (St Albans, NH) system running the test chambers to capture real-time behavioral events (including experimenter-delivered events like cues and reward delivery, and subject-generated inputs like lever presses), allowing perievent analysis of neural activity and LFP power.

### Surgery

#### Electrophysiological Recordings

A subset of subjects (n = 13 rats; 10 female, 3 male) underwent bilateral electrode implantation surgery (n= 7 ELS [6F/1M], 6 CTL [4F/2M]). Stereotaxic surgery was performed under isoflurane anesthesia (2–5%) using aseptic techniques. For each surgery, rats were secured in a stereotaxic apparatus (Kopf, Tijunga CA) using blunt earbars. Hair on the scalp was removed and the underlying skin scrubbed with two sets of alternating washes of Betadine scrub and 70% ethanol. Optical ointment (Vaseline) was applied gently to protect the eyes. A midline incision was made with a scalpel, and the scalp and underlying fascia retracted laterally with hemostats. A probe attached to the stereotaxic was used to measure the DV and ML deviations of Bregma and lambda; deviations of more than 0.1mm were adjusted until the head was level. Coordinates for array implants were generated from an atlas (Paxinos and Watson, 1996) for infralimbic cortex (AP, +2.7; ML, ±0.5), and then holes drilled using a dental burr over the location. At each of these insertion sites, the underlying dura was retracted to ensure the wires were inserted directly into brain tissue. In addition, holes were drilled for skull screws (typically three on each hemisphere) and the location of the ground wire (one each hemisphere).

Skull screws were inserted, after which each array was inserted (AP, +2.7; ML, ±0.5; DV, −5.0 mm), with the left inserted first. Each electrode consisted of two 8-wire electrode arrays (circular array surrounding an optical fiber; each wire consisting of a 50-μm dia Teflon-coated stainless-steel wire spaced 500 μm apart; NM Labs, Denison, TX). Arrays were lowered slowly (approximately 0.5mm/min) to the final recording location and fixed in place with dental acrylic. After this, the ground wire was wrapped around the posterior skull screw, then inserted into the brain at the ground wire hole. This was then repeated on the right side, with a specific mount used to ensure the two array connectors were spaced correctly for the headstage tethers that would be used later on the recording rigs. Rats received intramuscular injections of the antibiotic Baytril and the NSAID analgesic Meloxicam-SR at the end of surgery. Rats were given a 7-day post-surgery recovery period before conditioned suppression training began.

#### Perfusion and Histology

Following the final behavioral test, rats were deeply anesthetized using isoflurane 4% and transcardially perfused with 0.9% saline followed by 4% paraformaldehyde. Electrode placements were marked by passing current from a 9V battery through each electrode wire. Brains were postfixed in 4% paraformaldehyde for at least 12h, followed by 36-48h in 20% sucrose as a cryoprotectant, then stored at −80°C. Tissue was sectioned at 20 μm and mounted onto SuperFrost Plus slides (Fisher Scientific) using a cryostat at −20°C and imaged using a light microscope (Leica) to confirm electrode placement.

### Data Analysis

#### Behavioral Analysis

During Early Life, behavior was assessed for pup weight and juvenile social exploration. These data were analyzed with a between-subjects (unpaired) t-test using ELS and unstressed Controls as the factors of interest.

During Adulthood, behavior for Conditioned Suppression was measured by using a suppression ratio. Presses made during the 30-sec cue presentation (CS) was compared the 30-sec period immediately prior to the cue onset (BL). The suppression ratio for each stimulus type (i.e., CS+ or CS-) was calculated as:

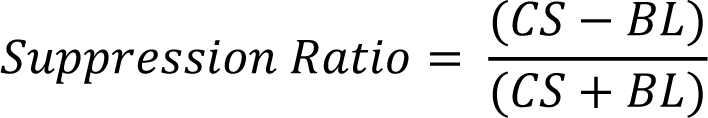

This produces a range of scores from −1 to +1, with −1 being total suppression of pressing during the cue and 0 reflecting no difference in pressing during the cue relative to the baseline. Differences between groups for behavioral suppression were determined using two-way ANOVA using the factors of Day (Days 1-3) and Stress (ELS vs unstressed Controls) as the variables of interest. Tukey’s HSD test was used for post hoc comparisons.

#### Single Unit Electrophysiological Analysis

Putative single units were sorted for each channel (wire) using principal component analysis clusters based on waveform similarity (Offline Sorter; Plexon). Unit clusters were then subject to secondary confirmation using auto-correlated firing properties. Auto-correlated firing histograms typically contain a “notch” at the 0 point indicative of a biologically-relevant refractory period for action potential generation (typically at least +/- 4 ms) Putative cells that showed significant numbers of spike events in this refractory period were rejected as units as being biologically implausible, and were not subsequently analyzed.

For perievent analysis, data were binned into 200ms blocks and averaged across events within a session. The perievent firing rate was then z-transformed based on mean and standard deviation of the average perievent activity. Thus, the z-normalized firing rate for each bin was calculated as:

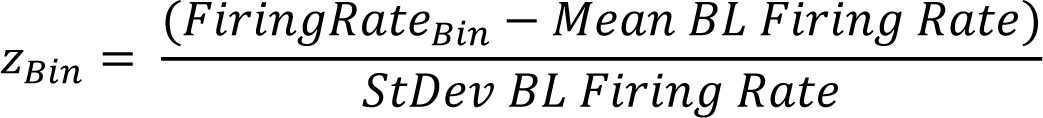

To ensure relatively uniform distributions of z-normalized firing, units with activity of less than 0.5Hz were excluded from subsequent analysis. Within the remaining units, averaged populations were grouped by generally excitatory (firing rate greater than 0.5z within 1s after cue onset) or generally inhibitory (firing less than −0.5z within 2s after cue onset). Based on this, we assessed two components of perievent cue firing. The first is the relative proportion of cells that exhibited generally excitatory (>0.5z), generally inhibitory (< −0.5z) or non-phasic relative to cue onset. These proportions were compared using chi square analysis. The second quantifies the peak firing during these onset periods for each cell. These were assessed using three-way ANOVAs with Stress (CTL vs ELS), Cue (CS+ vs CS-) and Epoch (Baseline vs Cue Onset) as factors.

#### Local Field Potential Analysis

LFP data generated spectrograms from 1-120 Hz perievent aligned to the CS+ and CS-cues. Spectrograms included a 5sec baseline followed by a 30s cue presentation and a 5-sec post-cue period, averaged into 200ms bins. Prior to fast Fourier transform (FFT), spectrograms were mean background subtracted, then normalized by the log of the Power Spectral Density (dB). From these spectrograms, specific frequencies were selected based on their established importance in circuit signaling: Delta (1-4 Hz), Low Theta (5-8 Hz), High Theta (9-14 Hz), Beta (15-22 Hz), Low Gamma (23-55 Hz), and High Gamma (65-95 Hz). The average power in these bands were then z-normalized by the average and standard deviation of the 5sec baseline period prior to cue onset, then applied to each 200ms bin throughout the perievent trace (similar to that described above for neural activity normalization).

For stress-related comparisons, for each spectrum, the subject’s baseline and average power during the cues (CS+ and CS-) was assessed separately for all days of the Conditioned Suppression tasks. Note that the averaged power during the cue period excluded the first 400ms of activity, but did include the rest of the cue period. These were assessed using three-way ANOVAs with Stress (CTL vs ELS), Cue (CS+ vs CS-) and Epoch (Baseline vs Cue Onset) as factors.

### Results Behavior

#### Early Life Stress Impairs Development

Pups were weighed six days following birth (PND6). We found that stress had a significant negative impact on weight gain, with animals in the ELS group showing reliably lower weights than Controls, *t*_24_ = 15.85, *p*< 0.0001 (Figure 1A).

**Figure 1.**
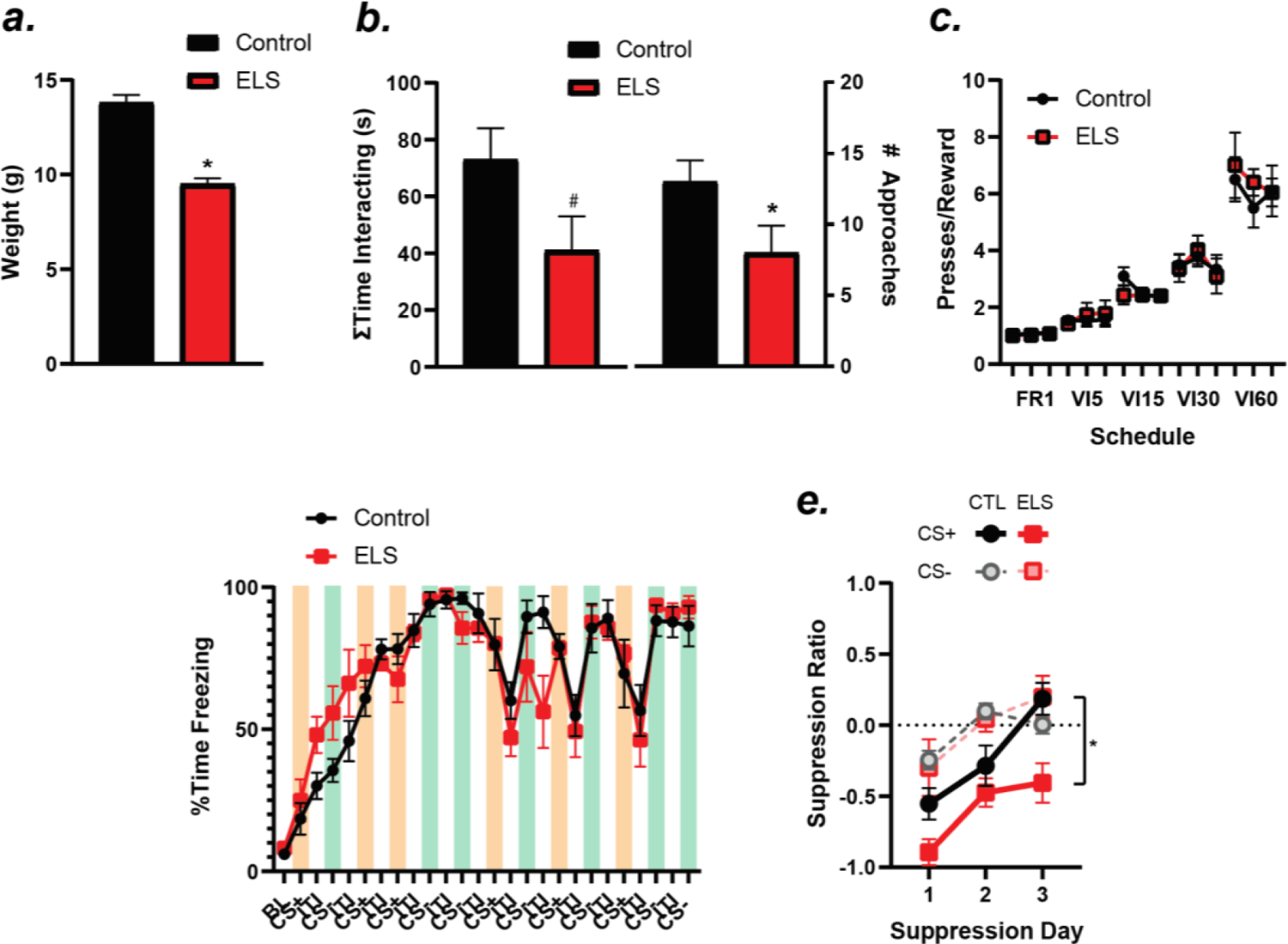
**A.** On PND6, weights of the ELS rats were lower than the unstressed Controls. **B.** In adulthood (PND180), ELS subjects showed less time (*left*) and fewer initiated contacts with a novel conspecific juvenile rat (*right*) in a JSI assessment. **C.** During Instrumental acquisition, rats in the ELS group showed similar levels of motivation to press as controls across decreasing schedules of reinforcement. **D.** Rats in both ELS and Control groups rapidly learned conditioned freezing to the presentations of CS+ (cue terminating in 0.8mA shock) and the “safe” CS-(no shock). Orange bars indicate freezing during the CS+ cue, green bars indicate freezing during the CS-. Data between colored bars indicate freezing during the intertrial interval between cues (i.e., contextual freezing). **E.** In a conditioned suppression task, the fear-associated CS+ cue suppressed lever pressing for food (VI60) more in ELS rats than Controls across three days of fear extinction. \**p*<0.05, main effect, ELS vs Control; #*p*<0.06, main effect, ELS vs Control; &*p*<0.05, ELS vs Control CS+ on that Day.

#### ELS Reduces Social Behaviors

After growing to adulthood in the vivarium (but prior to any experimental conditioning), adult rats were assessed in a juvenile social interaction (JSI) task on PND180 to assess social behaviors and anxiety-related phenotypes. Rats in the ELS group generally showed a decrease in social behaviors compared to Controls. While the total time spent sniffing the juvenile conspecific was marginally decreased in the ELS group (*t*_7_ = 2.32, *p*=0.052; Figure 1B, left), the number of interaction bouts initiated by the ELS subjects were significantly lower than unstressed Controls, *t*_7_ = 3.14, *p*=0.017; Figure 1B, right).

#### Instrumental Learning and Fear Conditioning are unaffected by ELS status

In the acquisition phase of instrumental learning, ELS appeared to have no effect on the motivation to press for food. Rats in the ELS group showed a similar ability as Controls on the last three days of each schedule to press for the food, and likewise to increase the number of presses per reward delivered based on the schedule requirements (Figure 1C). These observations produced a significant main effect of Schedule, *F*_2,223_ = 200.7, *p* < 0.001. This effect was almost exclusively due to a linear increase in press rate per reward earned across the decreasing reinforcement schedule, as a linear contrast on these data was significant, *F*_1,223_ = 274.1, *p* < 0.001 and accounted for 82% of the main effect variance. However, there were no effects of Stress or any interactions of Stress by any other factor (Schedule, Day) (all *F* < 1).

Following instrumental conditioning, rats learned fear conditioning to the CS+ and CS-tones in a novel context (Figure 1D). While both groups showed rapid acquisition of fear to the cues and context (main effect Trial, *F*_12,168_ = 48.72, *p* < 0.0001) and Cue (ITI vs Cue, *F*_1,14_ = 15.61, *p* = 0.001; greater freezing during cue), we did not see any Stress (*F*_1,14_ = 0.01, *p* < 0.91) or Stress X Cue X Trial interactions (*F*_12,168_ = 1.25, *p* = 0.25). Thus, there were no differences in the acquisition of conditioned fear between groups.

#### ELS Abnormally Suppresses Motivated Seeking Under Threat

Following this, rats were returned to the original context for the Conditioned Suppression task. Here, rats were reinforced on a VI60 schedule while receiving presentations of the CS+ (fear-associated cue; n=8) or neutral CS-(n=8). Because no shocks were delivered in this context, we repeated this Suppression paradigm for three consecutive days to assess the rate of fear extinction. For average pressing within each session, both groups showed suppression of lever presses during the presentation of the CS+, though ELS subjects were more suppressed than Controls (main effect Stress, *F*_1,16_ = 5.21, *p*=0.047; Figure 1E). This suppressive effect in the ELS animals was limited to the CS+ (interaction of Stress X Cue, *F*_1,16_ = 10.54, *p*=0.005), with ELS showing reliably greater suppression than controls during the CS+ (Tukey, *p*=0.005), but no differences between the CS-(Tukey, *p*=0.99).

Finally, we assessed the degree of successful extinction by assessing whether the suppression ratio was reliably negative (i.e., still suppressed) on each session. Controlling for multiple comparisons (Bonferroni), we found that in Controls, suppression for the CS+ was reliably below 0 on Day 1 (*p*<0.001), but not on Day 2 or Day 3. In contrast, ELS rats showed suppression during the CS+ that was reliably below 0 on all three days (Day 1: p<0.0001; Day 2: p=0.005; Day 3: p=0.01). Overall, these data indicate that relative to Controls, ELS animals display greater suppression of motivated behavior to fear-related stimuli that is more resistant to extinction than in Controls. However, ELS rats do not appear to show generalized fear, as they are adept at discriminating between fearful and “safe” stimuli; indeed, even better than Controls.

#### ELS increases the rate of excitatory responses to fear cues in vmPFC neurons

Recordings of single unit activity were conducted during Conditioned Suppression in both Controls (*n*=6) and ELS (*n*=7) subjects. From these recordings, we identified *n*=129 neural units in the Controls and *n*=191 in the ELS subjects in histologically-confirmed locations in the vmPFC based on boundaries found in (Paxinos and Watson, 1996) (Figure 2).

**Figure 2.**
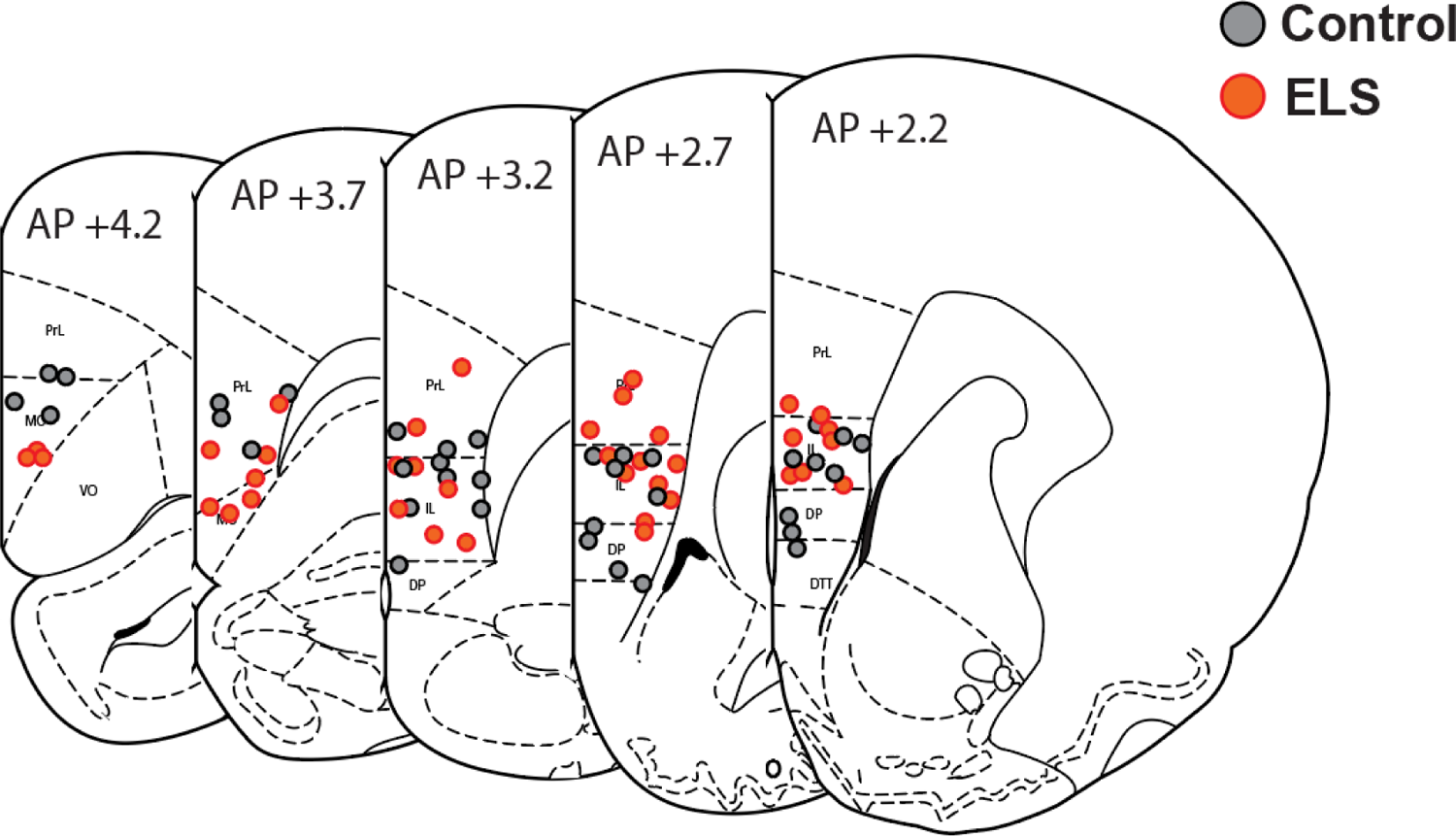
Placements of array wires in the PFC. Controls (black/gray circles) and ELS animals (red/orange circles) are shown primarily in the infralimbic cortex, with some wires extending ventrally into the medial orbital and dorsal peduncular cortex, and some dorsally into prelimbic cortex. Diagrams of brain and boundaries adapted from Paxinos & Watson, 1996.

Z-normalized firing rates were then aligned by their phasic response to the onset of the fear-associated cues (CS+ and CS-) by taking the average Z score during the first 1sec following cue onset. Data were considered generally excitatory if they exhibited an increase in firing greater than +0.5z relative to baseline, while inhibitions were those with a phasic response less than −0.5z. Data from all recorded cells relative to CS+ onset are shown in Figure 3A (Control) and Figure 3B (ELS). Population responses to the CS+ cue in the Controls were biased towards inhibitory signaling with 36.7% of cells demonstrating phasic inhibitions and 28.9% displaying excitations; 34.3% of cells were non-phasic in either direction. In contrast, ELS neurons showed the opposite pattern, as these units were almost twice as likely to display an excitatory response (50.5%) than an inhibitory response (25.5%) to the CS+ cue. ELS neurons also had slightly fewer cells that were non-phasic (24.0%). This shift from inhibitory to excitatory response to the CS+ between groups was significantly different, χ^2^ = 10.96, *p* = 0.0009 (Figure 3B). Notably, these EXC/INH/non proportions were quite stable by groups over days, with Controls showing generally more inhibitory responses than excitations to the CS+, and ELS showing the opposite pattern (*Controls*: Day 1 - 29% EXC, 35% INH; Day 2 – 32% EXC, 53% INH; Day 3 – 33% EXC, 27% INH; *ELS*:

**Figure 3.**
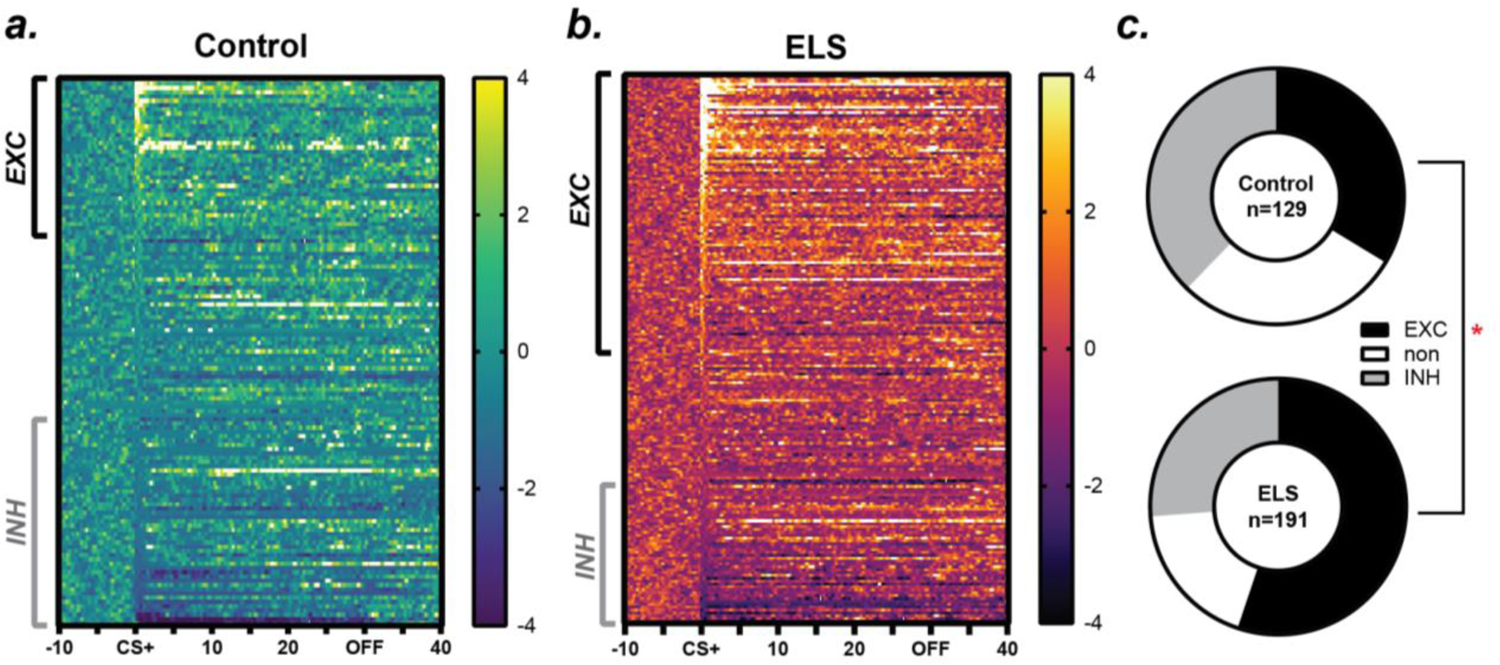
Heat plot representation of the population of recorded neural activity in the IL in Controls (**A**) and ELS subjects (**B**) relative to the onset of the CS+ cue in the Conditioned Suppression task. Color reflects magnitude of z-normalized firing with lighter colors indicating greater firing rates (z > 1), while darker colors indicate inhibitory activity (z < 1). Cells on the plot were sorted by the magnitude of the average firing rate during the first 1000ms after cue onset. Brackets on the left of each plot indicate the range of cells for which the phasic response was at least +0.5z above baseline (“excitatory”; *black top bracket*) or at least −0.5z below baseline (“inhibitory”; *gray bottom bracket*). Bar to the right of each heatplot indicates the scale to translate z score (from +4 to −4z) for each plot. **C.** Relative proportion of excitatory (EXC; greater than +0.5z), inhibitory (INH; less than −0.5z), and non-phasic units relative to the first 1sec of cue onset. ELS animals showed a significant increase in the proportion of EXC cells relative to Controls, χ^2^_1_ = 10.96, *p* = 0.0009

Day 1 - 56% EXC, 20% INH; Day 2 – 48% EXC, 31% INH; Day 3 – 40% EXC, 21% INH; all χ^2^ comparisons between day, p>0.20). In contrast to the fear-associated CS+, the relative proportion of excitatory and inhibitory response to the CS-cues was not different between groups (Control: 29.2% excitatory vs 32.1% inhibitory; ELS: 41.7% excitatory vs 34.9% inhibitory; χ^2^_1_ = 0.57, *p* = 0.45). These data suggest that ELS experience alters the function of the vmPFC to bias neurons towards an abnormally excitatory response to threatening (but not neutral, or “safe”) cues.

#### ELS impairs normal shifts in extinction-related firing to the CS+ cue

Prior investigations have reliably demonstrated that vmPFC neurons are critical for mediating extinction of fear via connectivity with amygdalar structures (Adhikari et al., 2015; Maren and Quirk, 2004; Milad and Quirk, 2002; Sierra-Mercado et al., 2011b). Phasic perievent excitatory activity relative to the CS+ in the Controls was consistent with this established finding, demonstrating a slight increase in the magnitude of phasic activity over days. In contrast, vmPFC neurons in ELS animals showed the opposite pattern, with the greatest level of excitatory activity during the CS+ occurring on the first day and decreasing magnitude of this response over repeated days of extinction (Figure 4A). In general, the ELS animals showed a reliably higher overall excitatory phasic response to the fear cues (main effect Stress, *F*_1,133_ = 4.77=8, *p* = 0.03) and an interaction of Cue (CS+ vs CS-) X Stress (ELS vs Control) X Extinction Day (1-3), *F*2,133 = 3.34, *p* = 0.04. This interaction showed that the phasic response to the CS+ was greater in ELS than Controls on Day 1 (*p* = 0.005) but not on subsequent days. There were no differences to the CS-between groups on any day (Figure 4B). However, in general, CS+ elicited significantly greater activity than the CS- in both the Controls (*p* = 0.02) and in the ELS group (*p* < 0.001).

**Figure 4.**
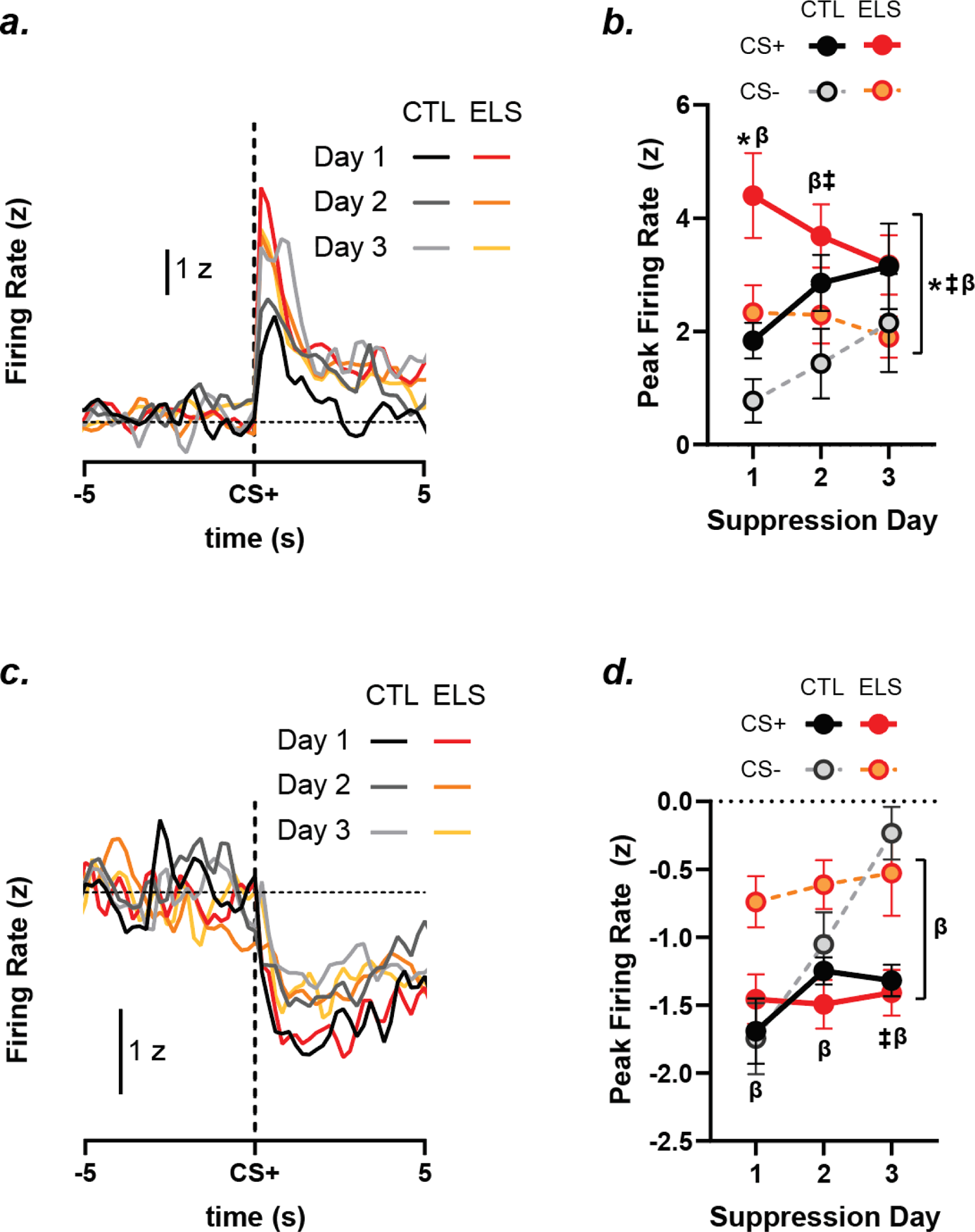
Phasic responses of vmPFC neurons to cue onset over repeated sessions of Conditioned Suppression. Identified excitatory (**A-B**) and inhibitory (**C-D**) units were analyzed separately. **A.** Both ELS (warm colors) and Control subjects (gray colors) showed rapid phasic responses to presentations of the fear-associated CS+ that typically lasted less than 1sec following cue onset. **B**. The average firing rate during the first 1sec following cue onset for each EXC cell for the CS+ (solid line) and the CS-(dashed line). C-D. Same as for A-B, but for maximum inhibitions (lowest firing point). \**p*<0.05, Control vs ELS (CS+);^ǂ^*p*<0.05, CS+ vs CS-(Controls); ^β^*p*<0.05, CS+ vs CS-(ELS).

In contrast to the excitatory phasic responses, for inhibitory responses, there were no stress-related main effect, *F*_1,104_ = 3.07, *p* = 0.08, though there was an interaction of Stress X Day, *F*_2,104_ = 3.18, *p* = 0.046, and Stress X Cue, *F*_1,104_ = 5.90, *p* = 0.017, but not Stress X Cue X Day, *F*_2,104_ = 0.75, *p* = 0.47 (Figure 4C). Indeed, for planned comparisons, we found no differences in the magnitude of the inhibitory response between Controls and ELS for the CS+ cues (all *p* > 0.28), though there were differences between ELS and Controls for the “safe” CS- on Day 1 (*p* < 0.001) and on Day 2 (*p* = 0.04), but not on Day 3 (*p* = 0.75). On those same days, ELS animals showed a better ability to discriminate neural firing responses between the CS+ and CS- on Day 1 (*p* = 0.005), Day 2 (*p* = 0.001) and Day 3 (*p <* 0.001), whereas Controls only successfully discriminated between CS+ and CS-cues on Day 3 (p = 0.01) but not on Day 1 (*p* = 0.75) or Day 2 (*p* = 0.76), Figure 4D.

#### ELS abolishes High Gamma LFP responses to threat cues in vmPFC

LFPs have been proposed to reflect the coherence of the aggregate voltage in a region. Given the large amount of surface area on dendritic arbors in a region relative to somatic activity, one potential interpretation of LFP oscillations is that it reflects a significant component of input to those local arbors from afferent regions. We recorded LFPs on the same wires and locations in the vmPFC as for the single-unit activity described above, and generated perievent spectrographs for defined frequency bands relative to CS+ and CS-onset during the same Conditioned Suppression sessions (West et al., 2021).

We found that ELS experience had little effect on changes in LFPs in most spectra (Figure 5). For example, in the Delta, Beta and Low Gamma frequencies, the response to the cue onset reliably decreased the power of these frequencies relative to baseline for the CS+ but not CS-(Cue [CS+ vs CS-] X Onset [baseline vs cue periods]: Delta, *F*_1,53_ = 14.52, *p* = 0.0004; Beta, *F*_1,53_ = 8.90, *p* = 0.004; Low Gamma, *F*_1,53_ = 11.66, *p* = 0.001). However, while the LFP response to the CS+ decreased LFP power in these spectra below baseline for the CS+ (all p<0.003) but not CS-, there were no differences between Controls or ELS for either CS+ or CS- in any of these spectra (all *p* > 0.25). Notably, there were no main effects of Stress or interactions of Stress with other factors in the Delta, Low Theta, High Theta, Beta, or Low Gamma frequencies.

**Figure 5.**
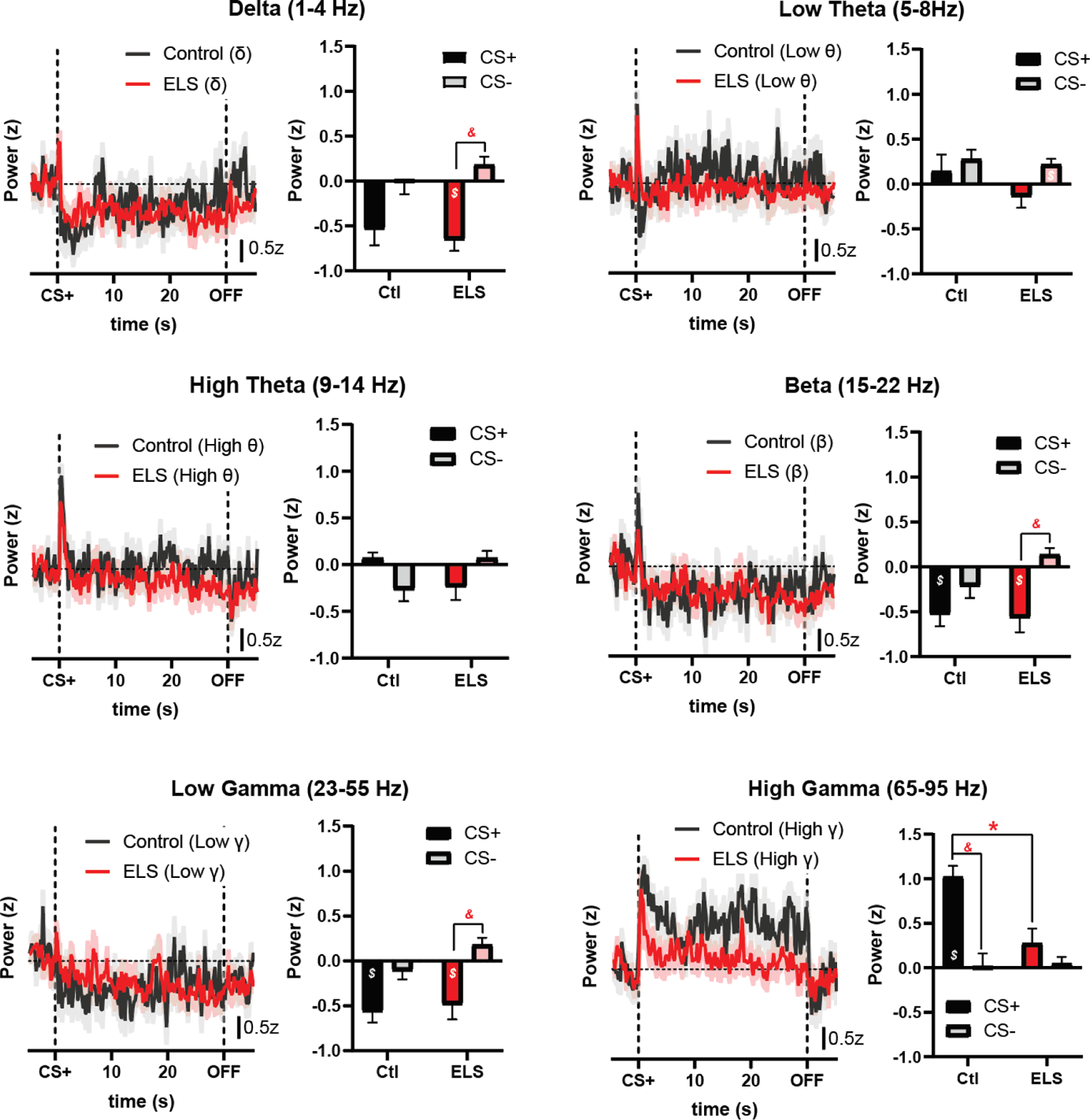
Perievent spectrograms generated for each of the frequencies identified in each title. Data are z-normalized by the average power in the baseline for each subject. At left in each subfigure is the mean response in 200ms bins over the duration of the CS+ cue presentations (Control: *black*; ELS: *red*). Vertical dotted line indicate cue onset and offset respectively. At right in each subfigure is the average (excluding the first 400ms, which may reflect a non-associative artifact). At left in black/gray are controls, and at right in red/pink are the ELS averages. In each pair, the darker/left bar is the CS+, while the lighter/right bar is the CS-. *p<0.05, Control v ELS; ^$^p<0.05, Baseline period vs Cue period; ^&^p<0.05, CS+ vs CS-.

However, ELS appeared to selectively impair the High Gamma frequency response (Figure 5, *bottom right*). In Controls, this response was a large and sustained increase in power for the duration of the CS+ cue, which then returned to baseline after cue offset. In contrast, this frequency response showed only a brief <1sec response to cue onset before rapidly returning to baseline for the rest of the CS+ cue. Quantifying the mean response during the cue period for both CS+ and CS-(Cue) for ELS and Controls (Stress) for the baseline period vs cue period (Onset), an ANOVA found a main effect of Stress, *F*_1,53_ = 6.89, *p* = 0.011, and an interaction of Stress X Cue, *F*_1,53_ = 6.76, *p* = 0.012, Stress X Onset, *F*_1,53_ = 6.90, *p* = 0.011, and Stress X Cue X Onset, *F*_1,53_ = 6.76, *p* = 0.012. Posthoc comparisons of this 3-way interaction indicated that in Controls, the High Gamma response to the CS+ was reliably higher than both its own baseline (*p*=0.0001) and the CS-cue (*p*=0.0001). The power for the CS-cue in Controls was, however, no different than its preceding baseline (*p*=0.99).

For ELS animals, this selective CS+ related increase in power was eliminated; the average power of the CS+ was no different from baseline (*p*=0.40) nor from the CS-period (*p*=0.99). Consistent with this loss of High Gamma power in the ELS group, there was a significant decrease for the High Gamma band for the CS+ cue between groups (*p*=0.0008), but there were no differences in power between groups for the CS-cue (*p*=0.99). As such, ELS appears to selectively abolish a discrete component of the LFP spectra, while leaving other lower-frequency components relatively unaffected.

## Discussion

Fear is an adaptive response to potentially threatening stimuli, though the brain must be adaptive enough that fear can extinguish when threats are no longer present. Consistent with prior observations, we found that ELS experience increases the fear-related suppression of reward seeking during the fear-associated CS+ compared to Controls. However, ELS animals did not differ from controls in their responses to the “safe” CS-cue. During these conditioned suppression sessions, recordings were made in the vmPFC that permitted the recording of both single unit and LFP activity in the same location. Neurons in the vmPFC of ELS rats showed both an increase in the proportion of excitatory responses to the fear-associated CS+ cue compared to Controls, as well as an increase in overall magnitude of the excitatory phasic response to cue. In contrast, while there were no differences between ELS and Controls in inhibitory encoding of the CS+, ELS neurons were better able to discriminate between CS+ and CS-stimuli than those in Controls. Finally, LFP oscillations in the vmPFC were consistent with a selective loss of the high gamma band in ELS-experienced rats. This loss is notable in that this is the only frequency where Controls showed a phasic change in activity that discriminated between CS+ and CS-stimuli, suggesting this signal plays a potentially important role in facilitating fear discrimination and feedback during extinction. Collectively, these data are among the first to demonstrate ELS-related functional alterations in vmPFC activity and resultant changes in fear-related behavior.

In general, our finding in Controls are congruent with prior findings in neurotypical adult rats undergoing fear conditioning and extinction. Controls showed initial fear to the CS+ stimulus that resulted in a robust cessation of motivated pressing for food. However, these fear-suppressed behaviors rapidly returned to pre-suppression levels by the second day of extinction. In these same Control subjects, vmPFC activity showed an appreciable increase in phasic excitatory activity in response to the CS+ commensurate with the resumption in motivated seeking behavior and extinction of the fear-related suppression, while showing reliably less activity to the safer CS-cue. These data are consistent with prior work demonstrating the role of IL and the vmPFC in mediating extinction through new learning (i.e., that the CS+ is now associated with no-shock), and increases in excitatory activity in these regions to permit this plasticity. Prior work, for example, has shown that excitatory stimulation of IL via electrical current or channelrhodopsin is sufficient to expedite fear suppression and extinction (Adhikari et al., 2015; Giustino and Maren, 2015; Milad and Quirk, 2002), and which persists in subsequent days without the stimulation present.

The ELS animals showed a different pattern of results; unlike Controls, ELS animals showed greater overall amounts of fear suppression, while at the same time showing increased levels of excitatory responding in single units. It is essential to note that ELS animals face developmental alterations compared to unstressed neurotypical controls. For example, PFC regions continue to develop and integrate neuromodulatory afferents for several days (at least PND16) after birth, including mesocortical dopaminergic wiring and integration with amygdalar nuclei (Cunningham et al., 2008; Kalsbeek et al., 1988; Kroon et al., 2019; Yuan et al., 2021), producing lasting changes in excitability and functional properties of these networks (Muhammad et al., 2012; Zhang, 2004). Thus, for these ELS animals whose stress experience happened during this critical developmental window, functional responses of these neurons may not mirror those in neurotypical individuals. Indeed, the robust and consistent increase in excitability in these neurons suggests that for ELS animals, the typical relationship between greater excitability and faster extinction seen in neurotypical controls no longer holds.

These observations argue against an interpretation of ELS inducing a hypofrontal state where extinction of threats are unable to be extinguished by descending prefrontal networks. This outcome would be consistent with some prior work showing decreased activity in human populations during reward and risk processing (Birn et al., 2017), and frankly with our *a priori* predictions for this study. However, the increased excitability suggests instead that vmPFC neurons are appropriately responding to the threat posed by the CS+ cue and are appropriately increasing activity to drive extinction, but that this activity is not sufficient to dampen fear-induced suppression as in controls. A possible interpretation for this set of results is that extinction is a process that requires both new learning (CS+ no longer predicts threat) as well as feedback to stamp in those new associations. Recent findings and models are consistent with the importance of these potential feedback mechanisms to the PFC in normal fear learning and extinction (McNally et al., 2011). For example, during processing of fear stimuli, mesocortical dopaminergic input to the PFC (Vander Weele et al., 2018) as well as amygdalar input (Burgos-Robles et al., 2017) provide event-related information about fear threats to prefrontal networks.

Indeed, recent work has demonstrated that pathway-specific inputs from intercalated neurons in the basolateral amygdala to discrete components of dorsal and ventral PFC may differentially regulate feedback to gate continued fear or its extinction (Hagihara et al., 2021). If this is the case, then persistent increases in fear-associated excitability in vmPFC of ELS animals may not be due to an inability to detect threats, but rather for a PFC-amygdala network to cooperatively use error-related feedback to update cues to a new and less-threatening state. If so, then evidence should exist that ELS animals are missing arising information that could be relevant for this learning.

Consistent with this interpretation, LFPs in the high gamma band were largely abolished in ELS compared to controls. LFPs reflect aggregate voltage in a region, and given the density of dendritic arbors relative to somas, these changes in voltages in a region may biased towards reflecting afferent inputs to a region, via depolarization and hyperpolarization of dendrites receiving those signals. Support for this perspective was recently provided in models of calcium transient activity with GCaMP sensors in the dorsal striatum (Legaria et al., 2021). Given this, one hypothesis consistent with our data is that vmPFC in ELS animals is lacking relevant feedback on the efficacy of extinction learning, and this information may be provided via gamma band oscillations.

This loss may be important for several reasons for interpreting our results. First, gamma oscillations have been thought to reflect in part the activity of GABAergic interneurons (Buzsáki and Wang, 2012; Cho et al., 2020; Sohal et al., 2009), and thus the ELS neurons displaying a heightened excitability in this study may reflect the loss of this GABAergic regulation.

Compellingly, BLA afferents preferentially target PFC GABA interneurons during early postnatal development (Cunningham et al., 2008), suggesting these pathways may be particularly vulnerable to insult during early life. Furthermore, disruption of this pathways during early life development appears to functionally alter and impair these arising BLA-PFC pathways, well-characterized dysfunction of this pathway in ELS individuals and animal models (Fan et al., 2014; Guadagno et al., 2018; Ishikawa et al., 2015; VanTieghem and Tottenham, 2018). Another potential source of input may arise from the hippocampus, which has likewise been implicated in fear-related changes in behavior in ELS-experienced animals (Reincke and Hanganu-Opatz, 2017). Consistent with this interpretation, high-gamma electrical stimulations in the fibria-fornix preferentially enhanced coordination between PFC and hippocampus, suggesting a likely route of communication on this frequency (Helbing and Angenstein, 2020). Future investigations will need to investigate these pathways, and in particular, why this frequency is uniquely disrupted while others are relatively unaffected.

Finally, recent work has focused not only on responses to threats, but also how animals come to learn about stimuli explicitly predictive of no-threat (i.e., safety). For our task, the CS-cue served as a neutral stimulus without consequence, but also signaled the explicit absence of any possibility of shock. This information about this safe cue appears to be reflected quite differentially in the vmPFC of ELS and Control subjects. In general, vmPFC neurons in Controls did surprisingly worse at discriminating between the CS+ and CS- than in ELS animals. While both ELS and Controls were adequate at discriminating between CS+ and CS-stimuli, this was not the case in inhibitory responses. Controls only showed discrimination between CS+ and CS- on the third day of fear extinction, while ELS animals showed robust and reliable discrimination between the CS+ and CS-throughout all days of extinction. These findings suggest the possibility that ELS animals may be more vigilant and ascribe a greater salience to potentially threatening stimuli, while therefore also better able to ascribe safety to non-threatening cues in the same context. This interpretation suggests that separate signals and neurons in the vmPFC may participate in the detection and significance of threat cues and their extinction (excitatory responses), while another participates in the learned safety of explicitly neutral stimuli (inhibitory responses). In this sense, ELS neurons were relatively impaired relative to Controls in excitatory signaling about threats and extinction, while they were relatively enhanced relative to controls in inhibitory signaling about safety signals. This intriguing dichotomy suggests discrete pathways that may coordinate complex responses to environments with ambiguous and competing information.

In conclusion, these data demonstrate that ELS is a potent modulator of brain networks that are essential for mediating appropriate and adaptive responses to a host of cognitive tasks including relief from fear, abstinence from drugs of abuse, and adequate assessment of risk in decision making. ELS experience, particularly early in development while the PFC and related limbic network are still in the process of developing functional connectivity, can have lasting effects on stimulus processing and behavioral responses to motivational stimuli. These data present new insights into how ELS-related dysfunction may contribute to the wide variety of mental health disorders that are precipitated by ELS and contribute to risk factors for disorders like addiction and PTSD.

### Limitations

The design of the ELS experience was not designed *a priori* as an early life stress model. As noted, the animals were originally destined for another project in our lab, and due to new animal care procedures in a new vivarium, the stressors that were presented were due to incidental damp bedding during a critical development period for both dams and pups. Prior work has demonstrated that unfortunate (but routine) vivarium stressors such as nearby construction can strongly alter animal behavior (Dallman et al., 1999). As such, we feel that these observations should be of interest not only to researchers interested in neural mechanisms of ELS, but also to veterinary and care staff who are interested in which disturbances in care environment can induce lasting changes in subject animals.

Second, we are aware that there are different and more standard ELS models of limited bedding, fragmented maternal care and others which have more extensive use in the field (Molet et al., 2014), although as noted, wet bedding as a stressor has some application in the field as well (Léonhardt et al., 2007). Understanding the conditions under more controlled experimental settings will provide a more comprehensive understanding on the impact of this ELS manipulation.

## Acknowledgements

This work was supported by a National Institute on Drug Abuse R00 Pathway to Independence award to MPS, along with a supplemental Research Supplements to Promote Diversity in Health-Related Research to MPS and FMB (DA035322). Thank you to Dr. Elizabeth West Niedringhaus for extensive help and answering all my questions about LFPs and also for providing input on an early draft of this document.

